# Dynamic healing and remodeling of mandibular ramus in segmental mobility and cortical overlap after intraoral vertical osteotomy

**DOI:** 10.1101/688978

**Authors:** Sang-Hwy Lee, Hoon Cho, Nam-Kyoo Kim, Bong Chul Kim, Jong-Ki Huh, Hyung Seog Yu, Hak-Jin Kim

## Abstract

The ramus immediately after intraoral vertical ramus osteotomy (IVRO) shows segmental mobility, cortical overlap, and projection, which yield smooth-surfaced continuity and stability after one year. This study aimed to elucidate the three-dimensional (3D) morphological changes driven by regional healing and remodeling 12 months postoperatively (***Post***) and their relationships with the immediate postoperative (***Imm***) segmental overlapping pattern. We performed a retrospective study of the morphological change of mandibular ramus in terms of time and amount of cortical overlap after IVRO. 3D computed tomography data of 108 hemi-mandibles were analyzed by independent superimposition of ***Imm*** proximal and distal segment to ***Post*** mandible. Analysis showed extensive regional bone resorption and apposition resulting in a relatively flat-surfaced and morphologically intact ***Post*** ramus. The middle ramal remodeling was significantly associated with ***Imm*** overlapping patterns and ***Post*** mandibular movement, new bony projections of various shapes being observed at the posterior border of the ***Post*** ramus. The ramus after IVRO underwent dynamic morphological changes during healing/remodeling which were partly associated with ***Imm*** overlap. Moreover, the ***Post*** functional and smooth-surfaced ramus seems to have been facilitated by the close surgical approximation of segments and the postoperative mandibular functional rehabilitation, even with segmental positional changes.

## Introduction

The intraoral vertical ramus osteotomy (IVRO), sometimes described as a vertical subcondylar or oblique osteotomy, is an orthognathic surgical technique for setting back the mandible[1, 2]. It is a common orthognathic procedure[3], along with sagittal split ramus osteotomy (SSRO), for prognathic or asymmetrical mandible. Though IVRO is advantageous as compared with SSRO[4, 5], certain factors must be taken into consideration for clinical application such as the induction of bone healing and remodeling[1, 6].

Since IVRO induces the distal segment to move backward, it overlaps with the proximal segment and becomes mobile to some extent without any intersegmental fixation. The bone healing commences in direct contact, with some mobility of cortical bones between two segments. Several studies have been conducted on the potential and outcomes of healing/remodeling, mainly based on histological findings[6-8], and on radiographic images[9-13]. However, these studies yielded limited insight into how the preoperative (***Pre***) ramus can be healed and remodeled to reshape the final postoperative (***Post***) one.

Over time, the remodeled ***Post*** ramus generally shows a different morphology from the original ***Pre*** and immediate postoperative (***Imm***) ramus, as seen in Fig 1. ***Post*** ramus generally presents reduced ramal height and width, reduced and smooth contoured angle, and different condylar angulation. The difficulty in understanding the transformation of ***Pre*** or ***Imm*** ramus to ***Post*** ramus may basically arise from the extensive morphological and positional change of the postsurgical segments. A few studies have reported good bone healing patterns based on three-dimensional (3D) analysis associated with IVRO[14], but without clear delineation of the IVRO ramus remodeling pattern[15-17].

**Fig 1.**
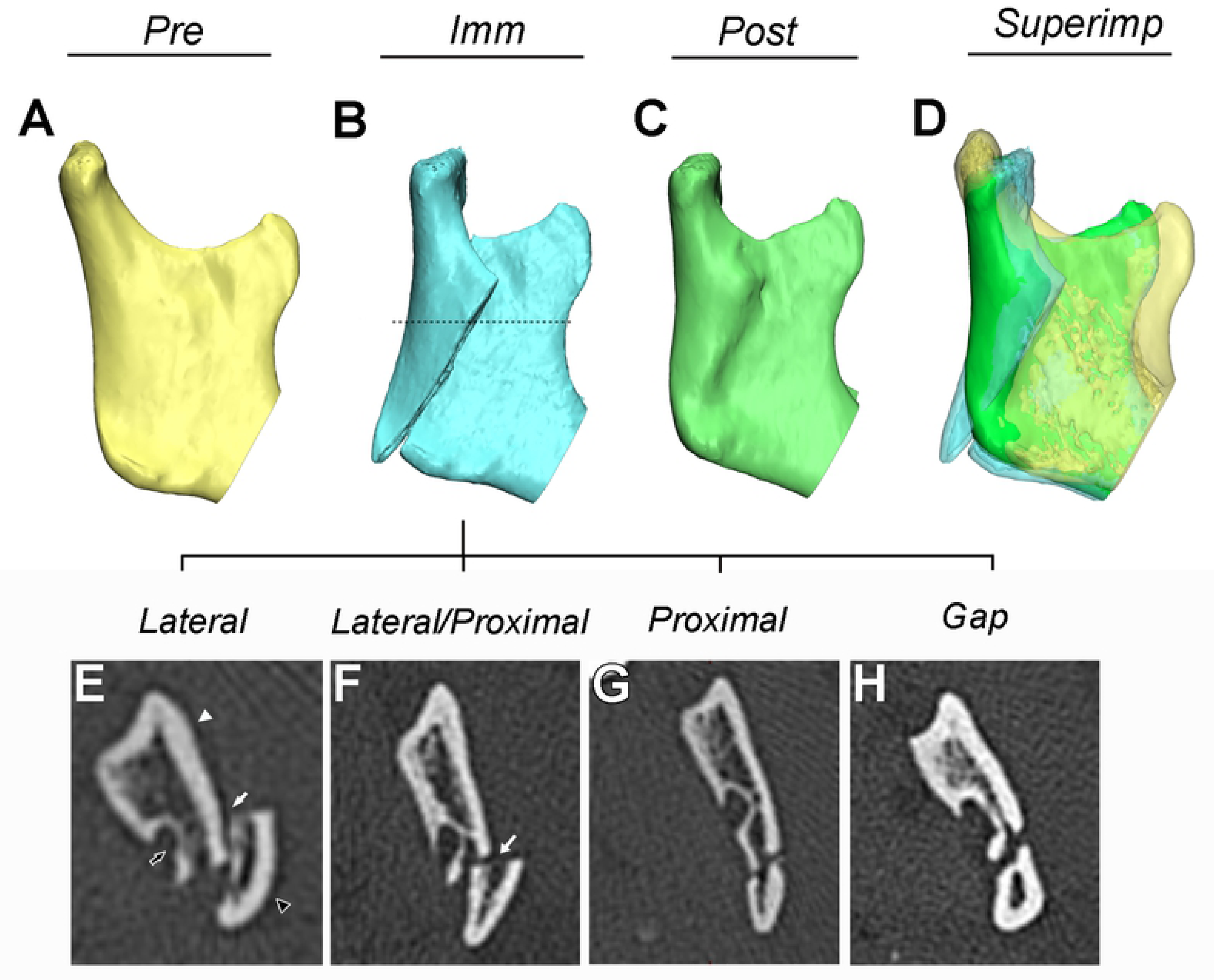
Morphological comparison of ramus at preoperative, immediate postoperative and one-year postoperative periods and its overlapping type between the proximal and distal segments at immediate postoperative stage after IVRO. The ramus was compared at three subsequent stages: (A) preoperative (***Pre***), (B) immediate postoperative (***Imm***), and(C) one-year postoperative (***Post***); (D) superimposition of the three stages. The ***Imm*** osteotomy gap was completely fused by bony healing and smoothened by remodeling on ***Post*** ramus. ***Post*** ramus also showed decreased ramal height and width, a smaller angular region, and a more upright and shorter condyle than ***Pre*** or ***Imm*** ramus.

The dotted line on the 3D model of ***Imm*** ramus in Fig B indicates the level of mandibular foramen entrance for judgments of overlapping pattern between the proximal and distal segments. (E) The positions of the proximal segment (black arrowhead) and the distal segment (white arrowhead) were evaluated on the axial images of CT data at the level of mandibular foramen (black arrow). The intersegmental line or space between the proximal and distal segment is indicated by a white arrow. (E-H) The ***Imm*** overlapping patterns between segments was classified into four types, mainly based on the position of the proximal segment in relation to the distal segment, as: (E) Lateral, (F) Lateral and Proximal, (G) Proximal and (H) Gap.

The primary purpose of this retrospective study was to investigate 3D morphological changes of ***Imm*** ramus as characterized by segmental mobility, cortical overlap, discontinuity, and projection after IVRO to stable ***Post*** ramus as seen in shape changes and smooth-surfaced continuity after one year. We hypothesized that the ***Imm*** ramus would be transformed into ***Post*** ramus via dynamic remodeling depending on elapsed time and mandibular function after surgery and also on the segmental overlapping pattern of ***Imm*** ramus. We also wanted to confirm that a direct model comparison of the ***Pre, Imm***, and ***Post*** rami would reveal interval morphological changes occurring in the ***Imm*** ramus attesting to regional bony remodeling. We introduced a novel direct segmental overlapping method to avoid potential bias induced by postoperative positional changes of the proximal/distal segments. The specific aims of this study were to: 1) verify the superimposition of fragmented ***Imm*** proximal and distal segments to ***Post*** ramus; 2) measure the inter-surface distance between ***Imm*** and ***Post*** ramus after IVRO to reveal the type and degree of surface remodeling; 3) determine the correlation between ***Post*** ramal remodeling and ***Imm*** segmental overlapping patterns or demographic variables; and 4) observe morphological changes and newly-formed structures after IVRO.

## Materials and methods

### Study Design and Sample

A retrospective cohort study was designed and implemented to address the research objectives. All subjects had presented for evaluation and management of malocclusion and facial deformity, and their inclusion criteria included those subjects who had unilateral or bilateral IVRO for mandibular prognathism and/or asymmetry and had undergone computed tomography (CT) examination with the same protocol at three time points, ***Pre, Imm*** and ***Post***. Subjects were excluded if they were previously operated, post-traumatic, tumor related, or syndromic. IVRO and postoperative management were performed for all subjects by the same protocol, including close approximation of segments by selective grinding, no intersegmental fixation, seven days of intermaxillary fixation, and six to eight weeks of staged active mandibular exercise[18, 19] for physiological bony healing and muscular rehabilitation. This work was approved by the local Ethics Committee of the Dental College Hospital, Yonsei University, Seoul, Korea (IRB 2-2015-0047).

## Study Variables

### Predictor Variable

The first variable was time after IVRO for healing and remodeling and the second was the intersegmental overlap pattern. To describe the second variable, the axial CT images of ***Imm*** rami were carefully evaluated at the level of the mandibular foramen entrance to evaluate patterns of overlap (Fig 1). These were classified into four types according to the position of the proximal segment in relation to the distal segment: ‘Lateral,’ ‘Lateral and Proximal,’ ‘Proximal,’ and ‘Gap’ (Fig 1E-H). This classification is derived and modified from previous reports[14, 20]. In addition, ***Post*** mandibular movement ranges during maximum interincial opening, protrusion, and lateral movement were considered the third variable.

### Outcome Variable

The first variable was the type and extent of surface remodeling, which included the bony apposition, resorption, and less marked change. The newly-formed bone surface of ***Post*** ramus as compared with that of ***Imm*** ramus was categorized as bony apposition and the reduced surface as bony resorption. The second outcome variable consisted of morphological changes of the ramus, including new structural formation.

### Data collection and analysis

CT scans were performed using a High-speed Advantage CT Scanner (GE Medical System, Milwaukee, U.S.A.) at Severance Hospital, Yonsei University Medical Center, Seoul, Korea. The imaging conditions were as follows: field of view 24.1 cm, 120 mA, 100 kV, scanning time 1 sec, and 0.5 mm thickness. The obtained CT data were reconstructed to produce 3D mandibular models for the three consequent stages of ***Pre, Imm***, and ***Post*** using Mimics (Version 17.0, Materialize, Leuven, Belgium) (Fig 2A, C, and E). The superimposition procedures for morphological comparison analyses were performed as follows (Fig 2):

**Fig 2.**
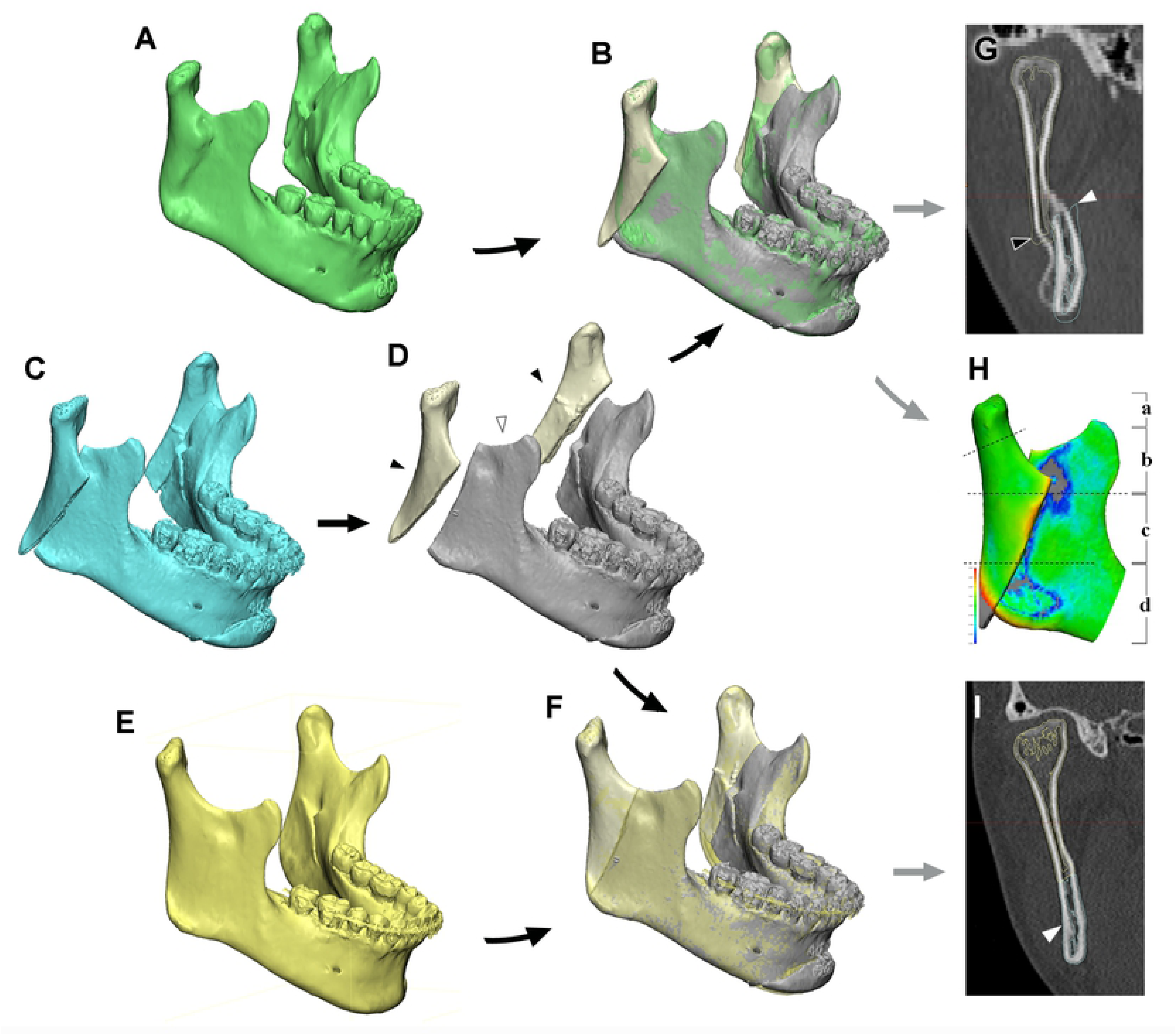
Workflow of the segmentation and superimposition with 3D mandibular models. The 3D mandibular models of one-year postoperative (***Post***) (A), immediate postoperative (***Imm***) (C), and preoperative (***Pre***) (E) stages were produced. ***Imm*** mandible (C) was divided into the proximal and distal segments at the osteotomy line, as seen in (D). These divided ***Imm*** proximal (indicated by black arrowheads) and distal segments (indicated by white arrowheads) were superimposed independently to the ***Post*** mandible (A), as shown in (B). The ***Post*** mandible was compared with the superimposed ***Imm*** proximal and distal segments by measuring the inter-surface distance (H) or comparing the outline (G) for evaluation of the ramal remodeling: 1) The comparison between ***Post*** and segmented ***Imm*** mandibles was made in four ramal regions, including the mandibular condyle (H-a), sigmoid notch (H-b), middle ramal region (H-c), and angle (H-d). 2) The inter-surface comparison between ***Post*** and segmented ***Imm*** mandible is shown in color maps (H), with the bony apposition in blue and the bony resorption in red. 3) The coronal CT image of ***Post*** ramus superimposed with the segmented and superimposed ***Imm*** ramus (G). The ***Imm*** proximal segment is indicated by a yellow line with black arrowhead, and the ***Imm*** distal segment as a blue line with white arrowhead. The separated ***Imm*** proximal and distal segments in (D) were also superimposed to ***Pre*** mandible (E) to verify the reliability of superimposition method, as shown in (F, H, and I). Note) proximal segment indicated by black arrowhead; distal segment by white arrowhead

The ***Imm*** mandible was segmented into proximal and distal segments at the IVRO osteotomy line (Fig 2D; white arrowhead for the segmented ***Imm*** distal segment; black arrowhead for the segmented proximal segment). The ***Imm*** distal segment was superimposed to ***Post*** mandible using two sequential steps of point-based and morphology-based registrations[21] (Fig 2B and G). The ***Imm*** proximal segment was also superimposed to ***Post*** mandible with the same independent procedure. The superimposed ***Post*** whole mandible and the proximal and distal segment of the ***Imm*** segmental mandible were compared to determine morphological and dimensional differences using 3-Matic (Version 10.0, Materialise, Leuven, Belgium) and Rapidform 2006 software (Inus Technology, Seoul, Korea) (Fig 2H). The differences between ***Post*** and ***Imm*** rami were measured at four ramal regions: the mandibular condyle (Fig 2H-a), sigmoid notch (Fig 2H-b), middle ramal region (Fig 2H-c), and angle (Fig 2H-d).

The data for the morphological descriptions and the inter-surface measurement values were collected. Descriptive statistics were computed for all study variables (R project, www.r-project.org). Fisher’s exact test was performed to find potential associations between predictor and outcome variables. In addition, analysis of variance (ANOVA) was made to evaluate the relationship between age and predictor or outcome variables. The level of significance was set at 0.05 for a 95% of confidence level. No subjects were lost for analysis in the study. The post hoc power analysis (G*Power 3.1.9.2, Heinrich-Heine-Universität Düsseldorf, Germany; PASS 15, NCSS, Kaysville, USA) was performed with an effect size of 0.4[22], an alpha value of 0.05, and total sample size of 108.

## Results

108 IVRO hemi-mandibles from 55 subjects met the inclusion criteria. The right and left ramus of each subject was independently analyzed after statistical correlation analysis by Chi-squared and Fisher’s exact probability tests for all outcome variables (p > 0.05; R project, www.r-project.org; details not shown).

The study consisted of 29 males and 26 females. Their ages at the time of surgery ranged between 17 and 41 years old (mean age 21.6 years old). Fisher’s statistical analysis (for gender) and ANOVA (for age) were performed to verify the associations between the gender or age and all the variables for ramal remodeling. No significant correlation was found between them (p > 0.05; Tables 1 and 2).

**Table 1.** The distribution of age and gender in relation to the overlapping types between proximal and distal segments at the immediate postoperative stage after IVRO.

| Overlapping | Lateral | Lateral/<br>Proximal | Proximal | Gap | Total | *p* |
| --- | --- | --- | --- | --- | --- | --- |
| Average Age | 21.9 | 21.0 | 21.4 | 20.6 | 21.6 | 0.697<sup>†</sup> |
| Male (sides) | 38 | 10 | 6 | 2 | 56 | 0.134<sup>‡</sup> |
| Female (sides) | 31 | 8 | 4 | 9 | 52 |  |
| Total (sides) | 69 | 18 | 10 | 11 | 108 | - |
| (%) | (63.9%) | (16.7%) | (9.3%) | (10.2%) | (100%) |  |

Note) <sup>†</sup>statistical analysis by ANOVA; <sup>‡</sup>by Fisher's exact test

**Table 2.** The distribution of age and gender in relation to the remodeling pattern at the lateral middle ramal area one year postoperatively after IVRO.

middle ramal area one year postoperatively after IVRO.
| Remodeling | Pronounced apposition/ resorption (> 2 mm) | Moderate apposition /resorption (1~2 mm) | Less apposition/resorption (<1 mm) | Total | *p* |
| --- | --- | --- | --- | --- | --- |
| Average Age | 21.8 | 21.3 | 20.9 | 21.6 | 0.580<sup>†</sup> |
| Male (sides) | 40 | 11 | 5 | 56 | 0.066<sup>‡</sup> |
| Female (sides) | 27 | 13 | 12 | 52 |  |
| Total (sides) | 67 | 24 | 17 | 108 | - |
| (%) | (62.0%) | (22.2%) | (15.7%) | (100%) |  |

Note) <sup>†</sup>statistical analysis by ANOVA; <sup>‡</sup>by Fisher's exact test

All subjects underwent mandibular setback (mean 6.7 mm, measured at the mandibular central incisal tip of the superimposed ***Pre*** and ***Imm*** 3D model) using bilateral IVRO (n=53/55) or unilateral IVRO (n=2/55) without any intersegmental fixation, simultaneously undergoing a Le Fort I maxillary osteotomy with semi-rigid fixation using miniplates.

### Predictor variables

The first predictor variable was the time lapse after IVRO for healing and remodeling. All subjects underwent CT examination before and after IVRO, at ***Pre*** (1-1.5 months preoperatively; mean 1.4 months), ***Imm*** (3-5 days postoperatively; mean 3.5 days), and ***Post*** (10 to 19 months). The mean time lapse for ***Post*** evaluation was 12.2 months after IVRO. ***Post*** outcome variables were not associated with the ***Post*** time lapse (Fisher’s exact test; p > 0.05; details not shown).

The second variable was the intersegmental overlap pattern during the ***Imm*** period, which was classified into four categories (Fig 1B, E-H; table 1). The laterally juxtaposed proximal segment in relation to the distal segment, classified as the ‘Lateral’ type (Fig 1E), was the most frequently observed pattern (N=69/108, 63.9%).

The third variable was the functional range of mandibular movement (Table 3). ***Post*** maximum opening for lateral overlapping group decreased 1.4-1.8 mm relative to that of preoperative period, but the amount of protrusion and lateral movement increased up to 2.6 mm.

**Table 3.**
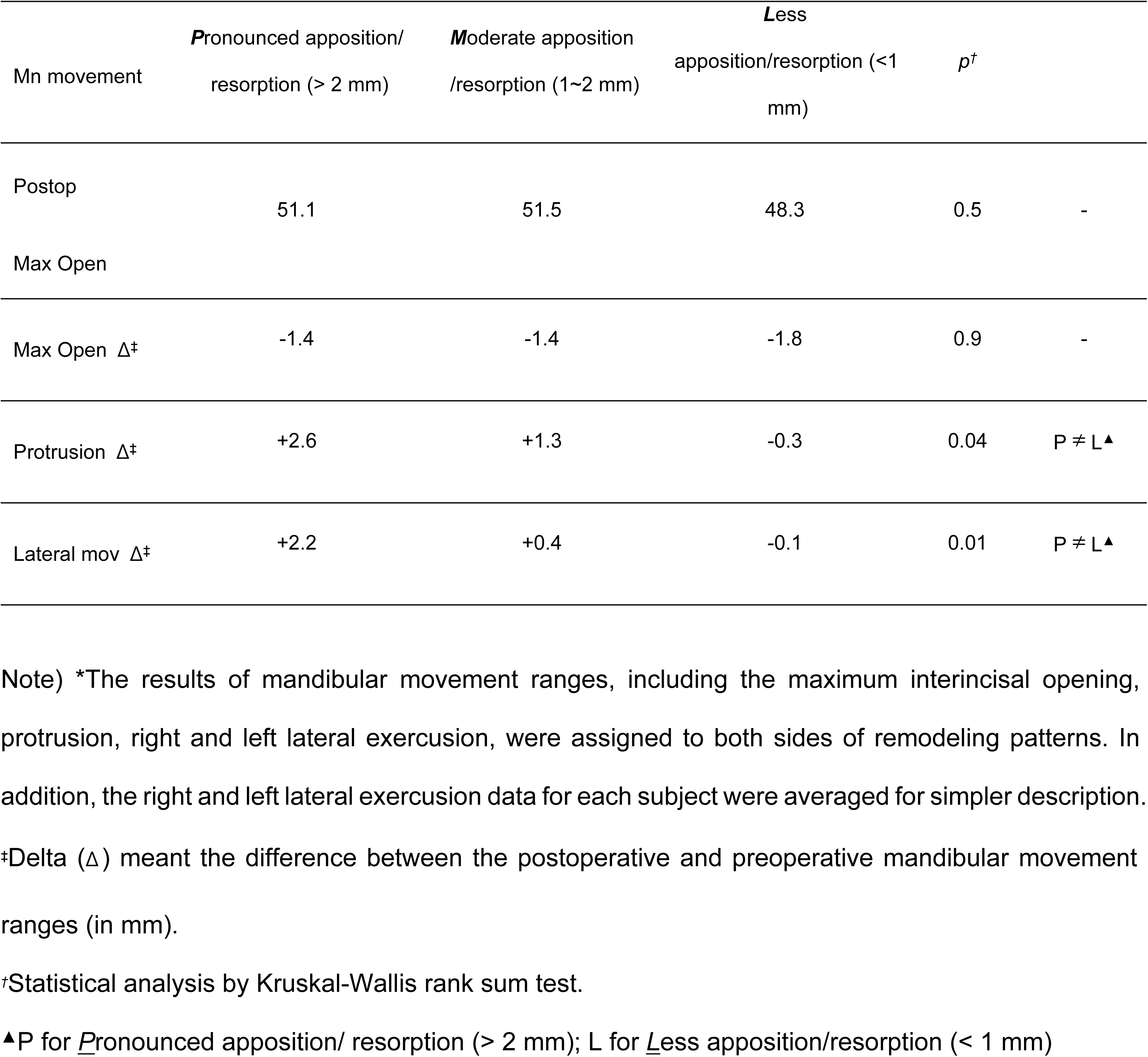
Comparison of mandibular movement ranges between pre- and postoperative period in relation to the remodeling pattern at the lateral middle ramal area for lateral overlapping group (total N= 69).*

### Outcome variables

The first outcome variable pertained to the type and extent of surface remodeling from ***Imm*** to ***Post*** ramus. These remodeling patterns were evaluated in four subdivisions of the ramus: the mandibular angle, sigmoid notch, middle ramus, and condylar region (as indicated in Fig 2H). Fisher’s statistical analysis was performed to confirm the association between the ***Imm*** segmental overlapping type (as a second primary predictor variable) and the ramal remodeling pattern (as an outcome variable) (table 3-6). In addition, the possibility of inter-regional correlation between these four subdivision regions were evaluated and rejected by the Chi-squared and Fisher’s exact tests (p > 0.05; details not shown). Details of remodeling in the four regions were as follows:

The most prominent bone resorption at the mandibular angle was found around the angular tip of the proximal segment on the lateral surface and the proximal tip of the distal segment on the medial surface (Fig 3E, F, K, L; indicated by black arrows; Table 4). The major bone apposition, on the other hand, was found at the angular tip of the distal segment on the lateral surface (Fig 3E, F, K, and L; indicated by white arrows).

**Table 4.** Remodeling at the mandibular angular and condylar region in relation to the intersegmental overlapping pattern.

| Regions | Remodeling Findings | Overlapping (N (%)) |  |  |  |  | P* |
| --- | --- | --- | --- | --- | --- | --- | --- |
|  |  | Lateral | Lat/Prox | Proximal | Gap | Total |  |
| Angular<br>Proximal | Pronounced resorption | 61 | 16 | 10 | 11 | 98 | 0.95 |
|  | (> 2 mm) | (85%) | (94%) | (100%) | (100%) | (89%) |  |
|  | Pronounced | 6 | 1 | 0 | 0 | 7 |  |
|  | apposition (> 2 mm) | (8%) | (6%) | (0%) | (0%) | (6%) |  |
|  | Less marked changes | 5 | 0 | 0 | 0 | 5 |  |
|  | (< 2 mm) | (7%) | (0%) | (0%) | (0%) | (4%) |  |
| Angular<br>Distal | Pronounced resorption | 32 | 5 | 0 | 3 | 40 | 0.15 |
|  | (> 2 mm) | (38.1%) | (26.3%) | (0%) | (27.3%) | (32.3%) |  |
|  | Pronounced | 43 | 11 | 7 | 6 | 67 |  |
|  | apposition (> 2 mm) | (51.2%) | (57.9%) | (70.0%) | (54.5%) | (54.0%) |  |
|  | Less marked changes | 9 | 3 | 3 | 2 | 17 |  |
|  | (< 2 mm) | (10.7%) | (15.8%) | (30.0%) | (18.2%) | (13.7%) |  |
| Condyle | Pronounced | 6 | 2 | 0 | 1 | 9 | 0.43 |
|  | apposition (> 2 mm) | (8.7%) | (11.1%) | (0%) | (9.1%) | (8.3%) |  |
|  | Moderate apposition | 27 | 6 | 1 | 5 | 39 |  |
|  | (1~2 mm) | (39.1%) | (33.3%) | (10.0%) | (45.5%) | (36.1%) |  |
|  | Minor apposition (< 1 | 36 | 10 | 9 | 5 | 60 |  |
|  | mm) | (52.2%) | (55.6%) | (90.0%) | (45.5%) | (55.6%) |  |

Note) \*statistical analysis by Fisher's exact test
Note) N-A for 'not available'; Lat/prox for lateral and proximal overlap pattern

**Fig 3.**
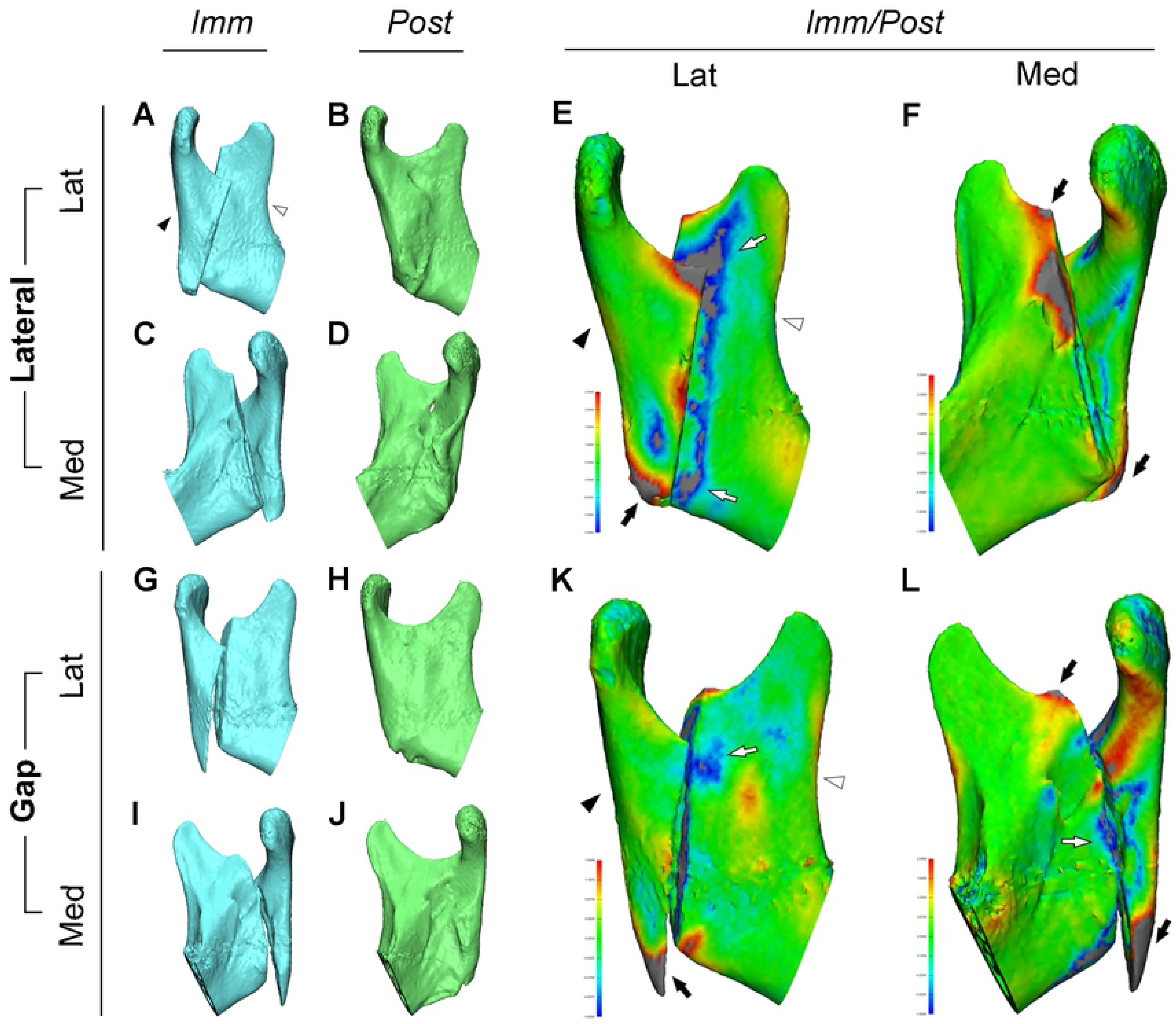
The ramal remodeling of ‘Lateral’ and ‘Gap’ overlapping types. (A-F) ***Imm*** ramus with the ‘Lateral’ overlapping type (A and C) was completely healed and remodeled, resulting in ***Post*** ramus (B and D). The ***Imm*** proximal (black arrowhead in A) and distal segments (white arrowhead in A) were independently superimposed to ***Post*** ramus (B and D) to reveal postoperative remodeling. The inter-surface distance between them was calculated and is shown in color maps for the lateral (E) and medial side (F) of ramus, indicating the high level of bony apposition (blue colored and indicated by white arrows) and resorption (red colored and indicated by black arrows). (G-L) The ‘Gap’ type ***Imm*** ramus (G and I) was also perfectly healed and remodeled to ***Post*** ramus (H and J). ***Imm*** proximal (blue in G and I) and distal segments (blue in G and I) were superimposed to ***Post*** ramus (H and J) to evaluate the postoperative remodeling with the resorption and bony filling at the intersegmental gap. The distance color maps between them are also shown on the lateral (K) and medial side (L) of ramus. Again, the bony apposition is shown in blue and indicated by white arrows and the resorption in red and indicated by black arrows. Note 1) Color coding for E and F; orange and red for 1.5 to 2.5 mm and gray for more than 2.5 mm of bone resorption; light and dark blue for −0.5 to −1.5 mm and in gray for more than −1.5 mm of bone apposition. Note 2) Color coding for K and L; in orange and red for 0.9 to 1.5 mm and in gray for more than 1.5mm of bone resorption; in light and dark blue for −0.4 to −1.0 mm and in gray for more than −1.0 mm of bone apposition. Note 3) ***Imm*** for immediate postoperative; ***Post*** for postoperative 1 year; ***Imm***/***Post***, superimposed comparison between ***Imm*** and ***Post***; Med, medial side; Lat, lateral side Note 4) proximal segment, indicated by black arrowhead; distal segment by white arrowhead; bone apposition, indicated by black arrows; and bone resorption, indicated by white arrows.

The middle ramal overlapping region between the proximal and distal segments also showed bony apposition and resorption at the lateral and medial surfaces, resulting in complete cortical continuity for all overlap types (Fig 3). The finding of ‘pronounced bony apposition/resorption’ prevailed in 67 cases (62.0%) on the lateral surface and 80 cases (74.1%) on the medial side (Table 5). The remodeling pattern in this region was significantly associated with the segmental overlapping type (Table 5; *p* < 0.001 for both lateral and medial regions). In addition, this remodeling was also significantly associated with ***Post*** mandibular movement ranges of protusion and lateral exercusion (Table 3; p < 0.04 and 0.01 each).

**Table 5.**
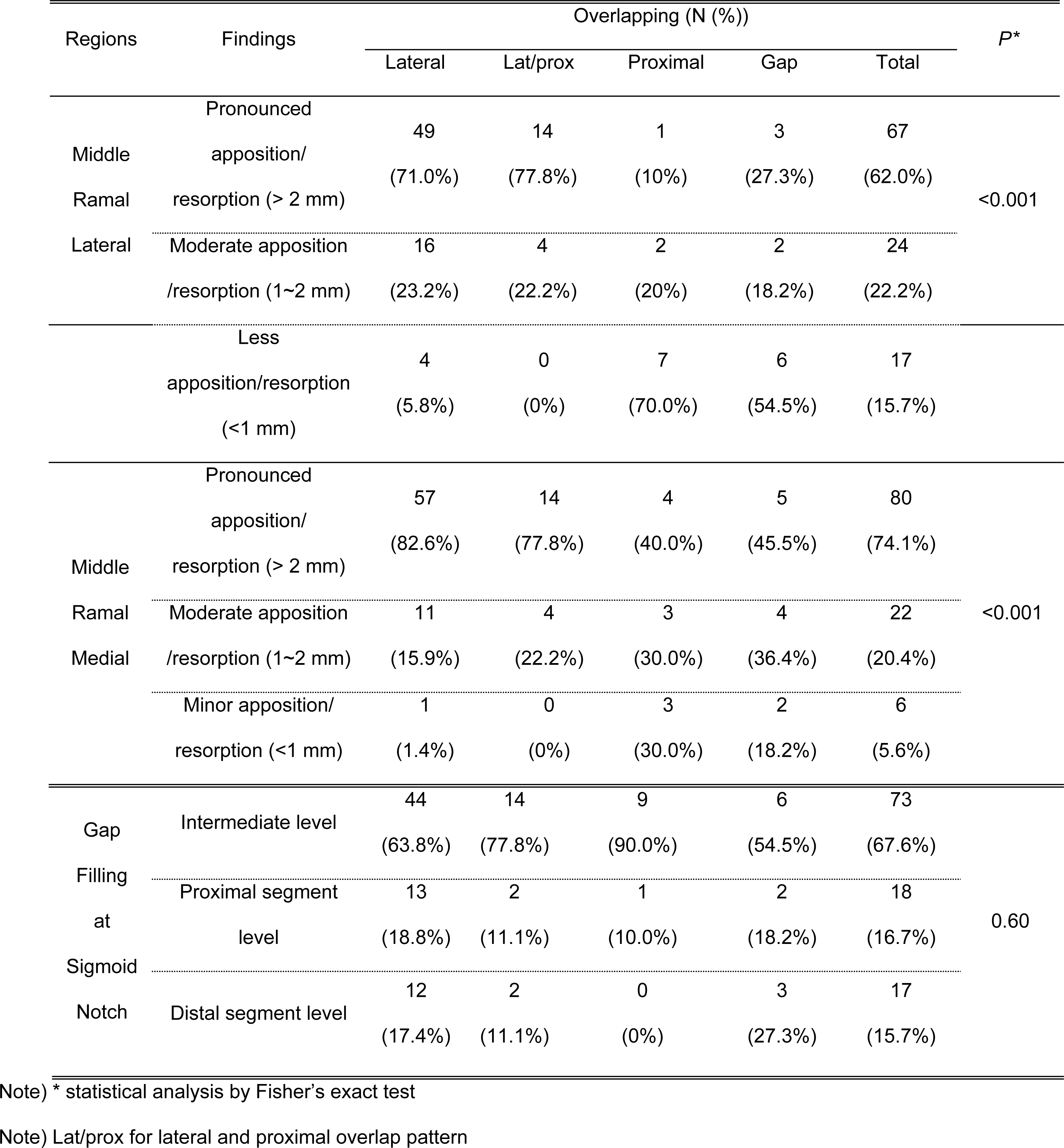
The remodeling pattern in the middle ramal and sigmoid notch region in relation to intersegmental overlapping type.

The sigmoid notch of ***Post*** ramus simultaneously showed both ‘moderate apposition and resorption’ of more than 1 mm in the majority of cases (88 cases, 81.5%), when compared with the ***Imm*** ramus (Fig 4; details not shown). The proximal segment near the osteotomy line mainly showed bone apposition (Fig 4B and D; indicated by black arrows), while the distal segment in the same region had resorption (indicated by white arrow).

**Fig 4.**
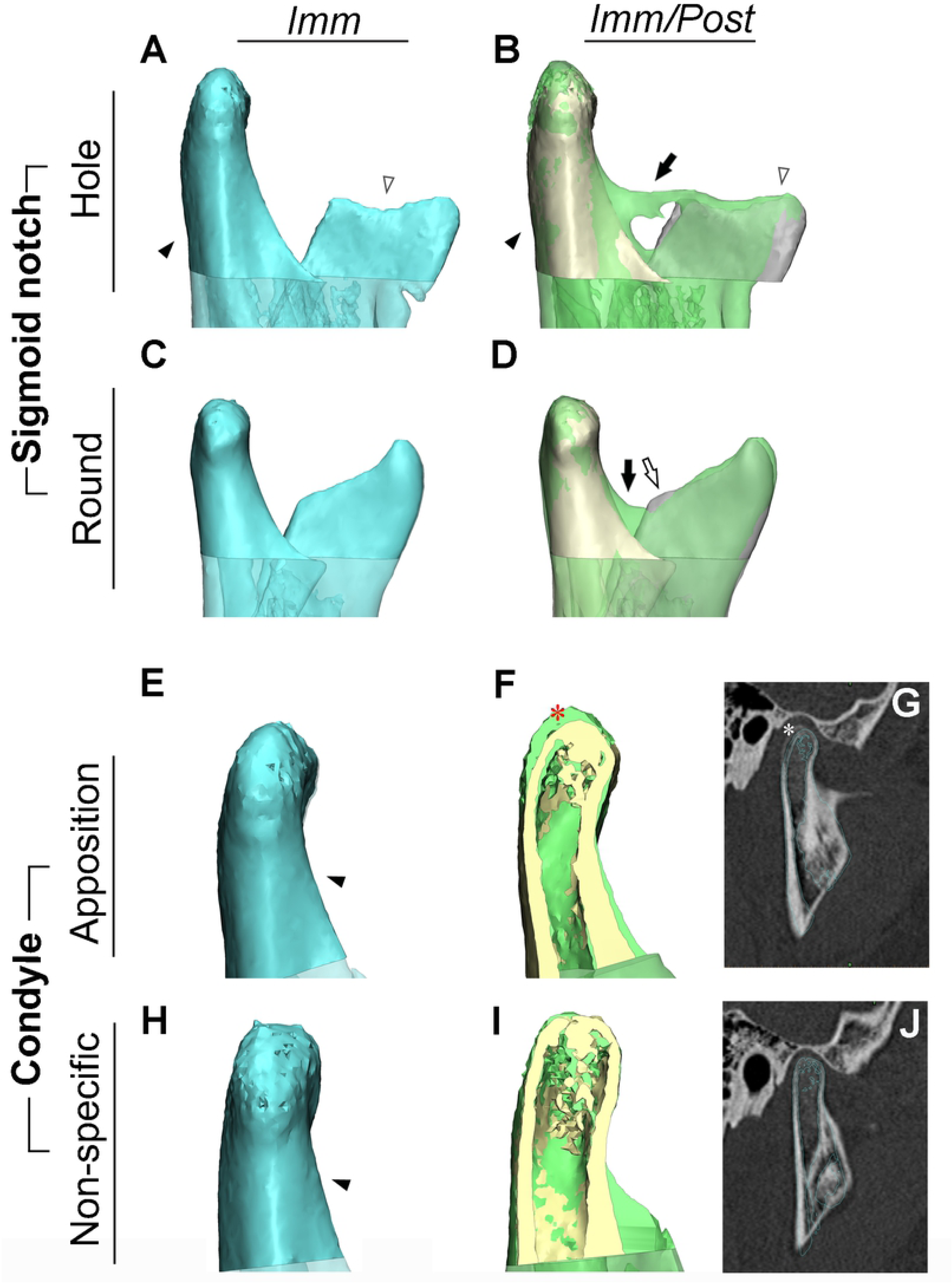
Morphological comparison of immediate postoperative (*Imm*) and one-year postoperative (*Post*) sigmoid notch and condyle. The sigmoid notch of ***Imm*** ramus shows the bony step between the proximal and distal segments (A and C), while the ***Post*** sigmoid notch is continuous in contour with different shapes (B and D). They were classified into hole (B), cleft, spicule, and round (D) type based on their morphological characteristics. (B and D) These show marked bony apposition and resorption. The ***Imm*** proximal (ivory, indicated by black arrowhead) and distal segments (gray, indicated by white arrowhead) were independently superimposed to ***Post*** ramus (light green) to reveal the ***Post*** remodeled ramus. The distance between them showed the level of bony apposition (indicated by black arrows) and resorption (indicated by white arrows). The condylar head of ***Imm*** and ***Post*** period did not reveal the definitive morphological or size differences (E and H for ***Imm***; F, G, I, and J for ***Post***). The ***Imm*** condyle (indicated by the black arrowhead in E and H, and ivory colored in F and I) was superimposed to ***Post*** condyle (light green colored in F and I) to reveal the ***Post*** remodeled ramus with different amounts of bone apposition. The 3D model and sagittal CT view from ***Post*** (G and J) also show the apposition of new bone (marked by asterisk in G) or none (in J). The calculated distance between them is shown in the color map, revealing the level of apposition. Note 1) ***Imm*** for immediate postoperative; ***Post*** for postoperative 1 year; ***Imm***/***Post*** (comparison of immediate and postoperative one year) for the superimposed comparison of ***Imm*** and ***Post*** ramus. Note 2) proximal segment, indicated by black arrowhead; distal segment by white arrowhead; bone apposition, indicated by black arrows; and bone resorption, indicated by white arrows; new bone, marked by asterisk (*).

The condylar region showed fewer marked bony changes than did other regions (Table 4 and Fig 4). 9 cases (8.3%) showed ‘marked apposition’ of more than 2 mm (Fig 4E-G), and 39 cases (36.1%) had ‘moderate apposition’ between 1 and 2 mm (Fig 4H-J). No notable finding of resorption was observed.

The second outcome variable consisted of morphological changes and new structure formation of the ramus. These were evaluated by Fisher’s statistical analysis to verify the association between the ***Imm*** segmental overlapping type and the ***Post*** ramal morphological characteristics. Details were as follows:

The sigmoid notch infrequently showed deformed patterns, including hole (Fig 4A and B), cleft, and spicule, while most cases had a normal round shape (64 cases, 57.1%; Fig 4C and D). Deformed patterns comprised hole-shape in 13 cases (12.0%) and cleft-shaped notch in 26 cases (24.1%), as shown in Table 6.

**Table 6.**
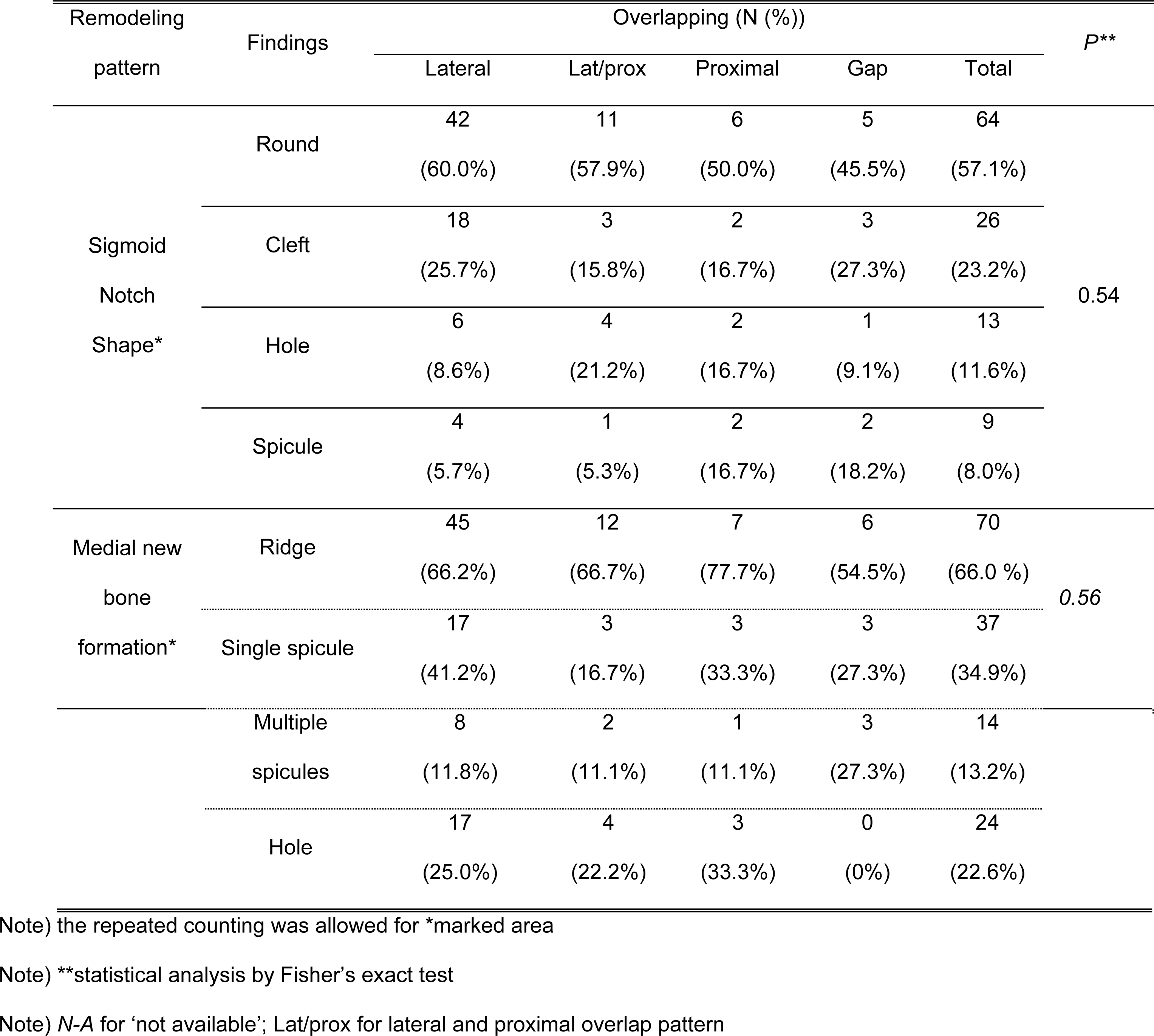
The morphological changes at the posterior border and sigmoid notch region in relation to intersegmental overlapping type.

The medial surface around the posterior border of ***Post*** ramus showed various shapes of new bone formation (Fig 5B, E, H, and K; indicated by black arrows). These were all so closely connected to the muscle that they seemed to be working as attachments of the medial pterygoid muscle (indicated by the dotted line in Fig 5C, F, I and L). These structures, not observed in the ***Imm*** mandible (Fig 5A, D, G, and J), were classified into four major types based on their morphological characteristics, including ridge, single spicule, multiple spicules and hole (Fig 5B, E, F, and K, respectively). 70 cases (64.8%) showed the ridge-shaped bony projection and 37 cases (34.3%) single spicule (Table 6). Their morphological types were statistically associated with the angular remodeling pattern (*p* < 0.046; details not shown), but not with the overlap pattern (p > 0.05; details not shown).

**Fig 5.**
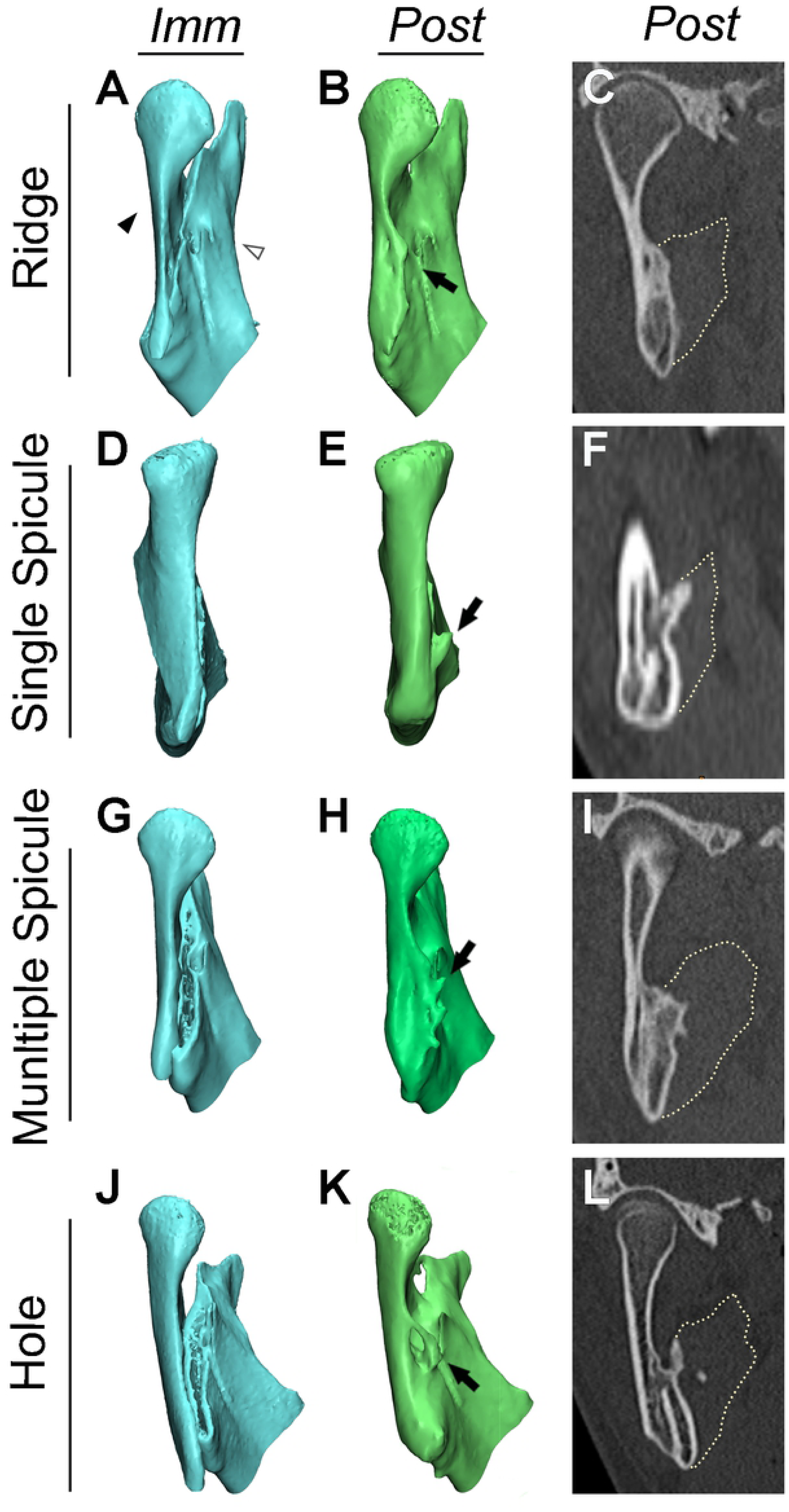
The new bone structures on medial surface of the posterior border. The ***Imm*** osteotomy gap between proximal (black arrowhead) and distal segments (white arrowhead) (A, D, G, and J) was completely healed and remodeled to ***Post*** ramus (B, E, H, and K). The posterior border of ***Imm*** ramus was remodeled to ***Post*** ramus with new bony structures (indicated by black arrows); these were classified into the ridge (B), single spicule (E), multiple spicules (H), or hole type (K), mainly based on their morphological characteristics. (C, F, I, and L) The dotted lines, indicating the outline of medial pterygoid muscle, were continuous to the new bone structures of medial ramus to suggest their relationships.

### Validation of methods

The reliability of the superimposition process was confirmed in three ways by two authors at one-week intervals. The first evaluation measured the mean distance in superimposed models between ***Imm*** proximal/distal segments and their corresponding ***Pre*** mandible (as outlined in Fig 2C-F), which was 0.0442 ± 0.0453 mm of inter-surface distance (N=6; intra-class correlation coefficient (ICC) between observers 0.990 and within observers 0.987 and 1.000 for observer 1 and 2). The second evaluation was performed by measuring the surface distance between the same sets of three ***Imm*** mandibles, which were superimposed independently to the same sets of ***Post*** mandibles (as shown in Fig 2A-D). Their mean distance was 0.0004 ± 0.0011 mm (ICC between observers 0.768 and within observers 0.812 and 0.924 for observer 1 and 2). The third validation, made during the superimposition procedures, verified coincidence of the cortical and marrow outline in the axial and coronal CT images (as seen in Fig 2G and I), though this was not quantitatively measured.

## Discussion

The primary purpose of this retrospective study was to investigate 3D morphological changes of the ***Imm*** ramus with segmental mobility and cortical overlap one year after IVRO. We hypothesized that ***Imm*** ramus was transformed into stable ***Post*** ramus via dynamic healing and remodeling in relation to the segmental overlapping pattern of ***Imm*** ramus. Our strategy, which employed a new direct segmental overlapping method, was to evaluate the remodeling pattern without being biased by the postoperative positional changes of the proximal and distal segments.

The ramal healing is one of the main concerns after IVRO in the case of intersegmental mobility without fixation. We expect the ***Imm*** ramus after IVRO to be healed by subperiosteal and endosteal calluses which fill the intersegmental gap and connect the separated segments[7]. Bell and Kennedy[6] reported that monkeys after 12 weeks of IVRO presented a continuous cortical structure of ramus with complete bony union and no evidence of medullary or cortical bone necrosis.

Ramal remodeling is another major concern after IVRO. Bone remodeling is a highly-coordinated cyclic process of mature bone tissue removal by osteoclasts and new bone formation by osteoblasts[23, 24]. Bone remodeling serves to adjust bone architecture to meet mechanical needs and/or to repair bone damage[25]. The primarily healed ramal structure undergoes remodeling through resorption, apposition, and rearrangement to meet functional and structural demands[6-10, 12, 13]. However, studies have shed little light on how the preoperative human ramus is remodeled to the postoperative one. Shepherd[11] and Westesson et al.[26] reported on morphological changes of ramus, describing postoperative angular resorption, decreased ramal width, increased cortical thickening, and decreased marrow space.

In our study, the overlapping region between the proximal and distal segments showed extensive bone formation and resorption on both the medial and lateral surfaces, contributing to the outcome of a flat-surfaced ramus (Fig 1 and 3 and Table 4). The distal tip of the proximal segment after setting back the distal segment was one of the most prominent regions, inviting extensive resorption of the ***Imm*** ramus, as reported previously[1, 7]. This remodeling pattern may be related to the protuberant high spot of the ***Imm*** ramus, decreased blood supply or bone necrosis[6], and the resultant reduction of ***Post*** height of proximal segment, though not correlated with the ***Imm*** overlapping pattern.

The range of mandibular movement, especially for protusion and lateral exercusion, was also confirmed as contributing factor to the formation of flat-surfaced ***Post*** ramus for the lateral overlapping group (Table 3). The skeletal muscle is functionally related to bone and the increased contraction of skeletal muscle affects the structure and biology of bone by increasing muscular vascularity, muscle mass and protein metabolism, and/or hormones/cytokines secreted by skeletal muscle during exercise[27, 28]. The importance of rehabilitation for muscle tissue and skeletal function after orthognathic surgery with systematic exercise regimen has been emphasized[29]. These observations therefore suggest the need for close approximation of the proximal and distal segments by selective grinding at the time of surgery and for ***Post*** active mandibular functional rehabilitation by systemic exercise to precipitate a smoother ramal surface.

Newly-formed structures were observed on the medial side of the posterior border in various shapes (Fig 5). These new bone formations may be related to the endosteal callus and the medial pterygoid muscle, since they were located just next to the osteotomy line and were in intimate contact with the medial pterygoid muscle on CT images (Fig 5C, F, I, and L). The proximal/distal segments are known to have endosteal callus after IVRO[1, 7], as well as the surgically detached and physiologically reattached medial pterygoid muscle[30].

Furthermore, these new structures were statistically associated with the angular remodeling pattern. We admit that the presence and shape of new structures may indicate the active rehabilitation of masticatory muscles, which in turn influences the extent or nature of remodeling in the angular region. These structures seem to be related to the physiological healing pattern of IVRO as well as to our treatment regimen. Also, we assume that the active masticatory muscle rehabilitation after IVRO helps the smooth-surfaced bony remodeling at the ramal overlap.

Our results are critically dependent on the accuracy of model superimposition. Generally the superimposition for 3D model comparison can be performed by rigid or non-rigid registration[31, 32]. To perform a direct 3D model comparison for ***Imm/Post*** ramus, we adopted a rigid surface-based registration method which utilizes the iterative closest point algorithm after the point-based preliminary registration. Our good error level and the matching of marrow/cortex indicated the strength of this superimposition method. However, the loss of positional information for the segments using this method inhibited the positional change-related remodeling analysis, though it could be compensated by the introduction of the overlap pattern.

Here we performed a direct 3D model comparative analysis of mandibular ramus one year after IVRO to observe the transformation of ***Imm*** ramus in terms of segmental mobility, cortical overlap, discontinuity, and projection to ***Post*** ramus with smooth-surfaced continuity and stability. This regional healing and remodeling was correlated with ***Imm*** segmental overlapping pattern and the range of mandibular movement at the middle ramus region. Moreover, the new bony projections with muscular attachments observed at the posterior border were correlated with the angular remodeling pattern. Further evaluation of 3D positional changes of the mandible and their relationship with the remodeling pattern is planned.

## Acknowledgments

This research was supported by a grant from the Korea Health Technology R&D Project, funded by the Ministry of Health & Welfare, Republic of Korea (grant number **HI17C0177**) for S.-H. Lee. We also want to express sincere thanks to Dr. Ji Wook Choi and Prof. Sang-Hoon Kang for their valuable contributions to this paper in the areas of sampling, modeling, analysis, and statistics.

## References

1. Boyne PJ. Osseous healing after oblique osteotomy of the mandibular ramus. J Oral Surg. 1966;24(2):125–33. PubMed PMID: 4955072.

2. Astrand P, Ridell A. Positional changes of the mandible and the upper and lower anterior teeth after oblique sliding osteotomy of the mandibular rami. A roentgen-cephalometric study of 55 patients. Scand J Plast Reconstr Surg. 1973;7(2):120–9. PubMed PMID: 4782117.

3. Ghali GE, Sikes Jr JW. Intraoral vertical ramus osteotomy as the preferred treatment for mandibular prognathism. J Oral Maxillofac Surg 2000;58(3):313–5.

4. Lee EM, Lee SH, Kim BC. Guided cutting of bone for intraoral vertical ramus osteotomy with a freer marking technique. Br J Oral Maxillofac Surg. 2015:660–1. Epub 2015/04/30. doi: 10.1016/j.bjoms.2015.04.001. PubMed PMID: 25921833.

5. Yu TH, Lim HJ, Lee J, Kim BC. Spontaneous Union of an Accidentally Fractured Proximal Segment During Vertical Ramus Osteotomy. J Craniofac Surg. 2017;28(4):1055–6. Epub 2017/05/27. doi: 10.1097/scs.0000000000003505. PubMed PMID: 28549042..

6. Bell WH, Kennedy JW, 3rd. Biological basis for vertical ramus osteotomies--a study of bone healing and revascularization in adult rhesus monkeys. J Oral Surg. 1976;34(3):215–24. PubMed PMID: 815526.

7. Lee SH, Park HS. Bone healing process in early mobilization after vertical ramus osteotomy of the mandible in adult dogs. J Korean Assoc Oral Maxillofac Surg. 1997;23(3):434–7.

8. Reitzik M. Cotrex-to-cortex healing after mandibular osteotomy. J Oral Maxillofac Surg. 1983;41:658–63.

9. Chung JH, Park HS, Hwang CJ. Morphologic and positional change of the proximal segments after intraoral vertical ramus osteotomy of the mandibular prognathism on submentovertex cephalogram. J Korean Assoc Oral Maxillofac Surg. 2003;29:26–34.

10. Pan JH, Lee JJ, Lin HY, Chen YJ, Jane Yao CC, Kok SH. Transverse and sagittal angulations of proximal segment after sagittal split and vertical ramus osteotomies and their influence on the stability of distal segment. J Formos Med Assoc. 2013;112(5):244–52. Epub 2013/05/11. doi: 10.1016/j.jfma.2012.02.013. PubMed PMID: 23660219..

11. Shepherd JP. Changes in the mandibular ramus following osteotomy - a long-term review. Br J Oral Surg. 1980;18:189–201.

12. Rhee BI, Park HS. On long-term remodeling of osteotomized segments after intraoral vertical ramus osteotomy in mandibular prognathism. J Korean Assoc Oral Maxillofac Surg. 1996;22(1):70–85.

13. Nihara J, Takeyama M, Takayama Y, Mutoh Y, Saito I. Postoperative changes in mandibular prognathism surgically treated by intraoral vertical ramus osteotomy. Int J Oral Maxillofac Surg. 2013;42(1):62–70. Epub 2012/08/04. doi: 10.1016/j.ijom.2012.06.024. PubMed PMID: 22858240..

14. Arimoto S, Hasegawa T, Kaneko K, Tateishi C, Furudoi S, Shibuya Y, et al. Observation of osseous healing after intraoral vertical ramus osteotomy: focus on computed tomography values. J Oral Maxillofac Surg. 2013;71(9):1602 e1–e10. doi: 10.1016/j.joms.2013.02.021. PubMed PMID: 23611606..

15. Ohba S, Nakao N, Awara K, Tobita T, Minamizato T, Kawasaki T, et al. The three-dimensional assessment of dynamic changes of the proximal segments after intraoral vertical ramus osteotomy. Cranio. 2015;33(4):276–84. Epub 2015/12/31. doi: 10.1080/08869634.2015.1097297. PubMed PMID: 26715130..

16. Ueki K, Hashiba Y, Marukawa K, Nakagawa K, Alam S, Okabe K, et al. The effects of changing position and angle of the proximal segment after intraoral vertical ramus osteotomy. Int J Oral Maxillofac Surg. 2009;38(10):1041–7. doi: 10.1016/j.ijom.2009.04.021. PubMed PMID: 19477622..

17. Katsumata A, Nojiri M, Fujishita M, Ariji Y, Ariji E, Langlais RP. Condylar head remodeling following mandibular setback osteotomy for prognathism: a comparative study of different imaging modalities. Oral Surg Oral Med Oral Pathol Oral Radiol Endod. 2006;101(4):505–14. doi: 10.1016/j.tripleo.2005.07.028. PubMed PMID: 16545716..

18. Fonseca RJ, Kenny JM. Oral and Maxillofacial Surgery: Elsevier; 2009.

19. Bell WH. Modern practice in orthognathic and reconstructive surgery: Saunders; 1992. 2517 p.

20. Kaneko K, Tateishi C, Imai Y, Hasegawa T, Fukuoka Y, Furudoi S, et al. Observation of Osseous Healing after Intraoral Vertical Ramus Osteotomy (IVRO). The Japanese Journal of Jaw Deformities. 2012;22(3):216–22. doi: 10.5927/jjjd.22.216.

21. Kim BC, Lee CE, Park W, Kang SH, Zhengguo P, Yi CK, et al. Integration accuracy of digital dental models and 3-dimensional computerized tomography images by sequential point-and surface-based markerless registration. Oral Surg Oral Med Oral Pathol Oral Radiol Endod. 2010;110(3):370–8. Epub 2010/07/02. doi: 10.1016/j.tripleo.2010.03.036. PubMed PMID: 20591700..

22. Cohen J. Statistical power analysis for the behavioral sciences. 2nd. : Lawrence Erlbaum Associate; 1988.

23. Raggatt LJ, Partridge NC. Cellular and molecular mechanisms of bone remodeling. J Biol Chem. 2010;285(33):25103–8. doi: 10.1074/jbc.R109.041087. PubMed PMID: 20501658; PubMed Central PMCID: PMCPMC2919071..

24. Crockett JC, Rogers MJ, Coxon FP, Hocking LJ, Helfrich MH. Bone remodelling at a glance. Journal of Cell Science. 2011;124(7):991–8. doi: 10.1242/jcs.063032.

25. Hadjidakis DJ, Androulakis, II. Bone remodeling. Ann N Y Acad Sci. 2006;1092:385–96. doi: 10.1196/annals.1365.035. PubMed PMID: 17308163..

26. Westesson PL, Dahlberg G, Hansson LG, Eriksson L, Ketonen L. Osseous and muscular changes after vertical ramus osteotomy. A magnetic resonance imaging study. Oral Surg Oral Med Oral Pathol. 1991;72(2):139–45. PubMed PMID: 1923390.

27. Salmons S. The adaptive response of skeletal muscle: What is the evidence? Muscle Nerve. 2018;57(4):531–41. Epub 2017/09/01. doi: 10.1002/mus.25949. PubMed PMID: 28857207..

28. Karsenty G, Mera P. Molecular bases of the crosstalk between bone and muscle. Bone. 2018;115:43–9.

29. Storum KA, Bell WH. The effect of physical rehabilitation on mandibular function after ramus osteotomies. J Oral Maxillofac Surg. 1986;44(2):94–9. PubMed PMID: 3456031.

30. Cruz DZ, Rodrigues L, Luz JG. Effects of detachment and repositioning of the medial pterygoid muscle on the growth of the maxilla and mandible of young rats. Acta Cirúrgica Brasileira. 2009;24(2):93–7.

31. Chen Y, Zhao J, Deng Q, Duan F. 3D craniofacial registration using thin-plate spline transform and cylindrical surface projection. PLoS One. 2017;12(10):e0185567. Epub 2017/10/06. doi: 10.1371/journal.pone.0185567. PubMed PMID: 28982117..

32. Cevidanes LH, Motta A, Proffit WR, Ackerman JL, Styner M. Cranial base superimposition for 3-dimensional evaluation of soft-tissue changes. Am J Orthod Dentofacial Orthop. 2010;137(4 Suppl):S120–9. Epub 2010/04/23. doi: 10.1016/j.ajodo.2009.04.021. PubMed PMID: 20381752; PubMed Central PMCID: PMCPMC2859472..

